# Variable and conserved features of copy-back viral genome populations generated *de novo* during Sendai virus infection

**DOI:** 10.1101/2025.10.21.683696

**Authors:** Yanling Yang, Yuchen Wang, Carolina B. López

## Abstract

Copy-back viral genomes (cbVGs) are generated during the replication of negative-sense RNA viruses when the polymerase drops off from the genome and reattaches to the nascent strand. cbVGs have strong immunostimulatory properties and impact infection outcomes. Despite their importance, the composition and mechanisms of *de novo* cbVG generation and accumulation remain unclear due to challenges in obtaining cbVG-free virus stocks (clean stocks). Here, we obtained several clean stocks by independently rescuing recombinant Sendai virus (SeV) six times and verified their cleanliness through PCR, RNA sequencing, and absence of immunostimulatory activity. High multiplicity-of-infection passaging of clean stocks produced six high-MOI passaged stocks, each with distinct cbVG populations. Among them, polymerase drop-off (break) positions occurred throughout the genome, while polymerase reattachment (rejoin) positions preferentially occurred near the trailer end. Few common breaks were observed between stocks, while there was a hot rejoin region near the trailer end. In each stock, a few cbVG species dominated and remained stable across passages, all conforming to the ‘rule of six’, regardless of length. Low-abundance cbVGs were variable across passages, indicating the continuous generation of new cbVGs, despite the stabilization of a subset of species. Intriguingly, cbVG species which originated from polymerase drop-off at or close to nucleotide 1 were present in all stocks, suggesting that cbVG species originating at the 3’ end of the genome are conserved products of SeV replication.

## Introduction

Copy-back viral genomes (cbVGs) are one of the most extensively studied classes of non-standard viral genomes (nsVGs) (1). Although initially considered artifacts that propagated along with standard viral genomes (stVGs), cbVGs are now recognized as key modulators of virus-host interaction during the course of infections (2). Moreover, the kinetics and dynamics of cbVG accumulation correlate with disease progression and clinical outcomes (3). The biological activity of cbVGs is accomplished through four major effects: interfering with standard virus replication, activating innate immune pathways via RIG-I and MDA5, promoting persistence through prosurvival signaling, and inducing cellular stress responses that suppress viral protein expression (4–6).

cbVGs are generated during the replication of negative-strand RNA viruses when the polymerase drops off (breaks) from the genome and reattaches (rejoins) to the nascent strand, using it as a template to complete replication (2, 7). Unlike standard genomes, cbVGs contain a unique junction region formed by the combined break and rejoin sites, as well as complementary trailer ends that are present only in cbVGs (2, 8). cbVGs have been observed in several mononegavirales, including Sendai virus (SeV), measles virus, parainfluenza virus 5 and respiratory syncytial virus (RSV) (9–12). SeV, especially the Cantell strain, has long served as a model for studying cbVGs as a single well-characterized cbVG, cbVG-546, has been consistently detected in SeV Cantell virus stocks and in samples from SeV Cantell infections (8). cbVG-546 is considered the prototypical immunostimulatory cbVG, activating RIG-I/MDA5 or TLR3 signaling, and its derivative, a defective viral genome-derived oligonucleotide (DDO), has been used as a vaccine adjuvant in pre-clinical studies (8, 13–19).

Beyond their direct applications, cbVGs play critical roles in viral replication and host responses, making the regulation of their generation an attractive target for therapeutic development (7). However, our understanding of viral and host factors that regulate cbVG generation and accumulation is limited to data showing that the SeV and measles virus C proteins suppress the production and accumulation of cbVGs (20–22), as well as a single amino acid substitution in the SeV N protein drives cbVG production (23). A key challenge in identifying drivers of cbVG generation is the difficulty of obtaining cbVG-free viral stocks, as cbVGs are often carried over from early isolates, leading to biased populations. This complicates efforts to distinguish *de novo* generation of cbVGs from the selection and accumulation of already existing species. For example, it remains unclear whether cbVG-546 in the SeV Cantell strain has been persistently maintained as the virus is passaged and expanded or whether it is newly generated during infection with cbVG-depleted SeV Cantell stocks (24). Resolving such questions is essential for a comprehensive understanding of cbVG biology and for effectively harnessing their therapeutic potential.

Towards this goal, we successfully generated multiple pure stocks of newly rescued recombinant SeV Cantell reporter virus (rSeVC^eGFP^) that were confirmed to be cbVG-free. By passaging these clean stocks at high multiplicity of infection (MOI), we obtained six distinct cbVG populations. Sequencing analysis revealed that break points occurred throughout the genome, with rejoin events preferentially occurring near the trailer end, and that every stock contained cbVGs with a break at position 1. Although diverse cbVG populations arose independently from the same virus, only a small subset of cbVG species was conserved across passages, and unexpectedly, shorter cbVGs did not preferentially accumulate as dominant species. Together, these findings reveal both diversity and reproducibility in cbVG generation and provide a foundation for defining the mechanisms that govern this process.

## Results

### Reverse genetics enables the generation of cbVG-clean Sendai virus (SeV) Cantell stocks

To obtain SeV stocks free of cbVGs, we independently rescued six recombinant rSeVC^eGFP^ virus stocks, which we named rS1-rS6. rSeVC^eGFP^ was designed based on the SeV Cantell strain, with eGFP inserted between the NP and P genes, as previously described (25) (Fig. 1A). Viruses were rescued by co-transfecting the full-length plasmid and three helper plasmids into BSR-T7 cells to generate passage 0 (P0) stocks. Passage 1 (P1) stocks were expanded from P0 by infecting LLC-MK2 cells at an MOI of 0.1 (Fig. 1B). To evaluate whether the P1 stocks were free of cbVGs, we used three different methods. First, we utilized a well-described PCR assay to amplify cbVGs present in the stocks (24). In this assay, both the stVG and cbVG are reverse transcribed using a single RT primer. The stVG is detected upon amplification of a segment of the virus with specific primers, while cbVGs are detected using the original RT primer and a primer complementary to the trailer of the virus. Because these two primers are oriented in the same direction on the stVG, no stVGs can be amplified (Figure S1). No obvious cbVG bands were detected in any of the six newly rescued P1 virus stocks infected cells, whereas the SeV genome was detected in all SeV stock-infected samples (Fig. 1C). Next, we used RNA sequencing followed by analysis with the Viral Opensource DVG Key Algorithm 2 (VODKA2) to detect cbVGs in each stock by precisely detecting their junctions (Break position_Rejoin position) with high sensitivity and specificity (26). VODKA2 analysis revealed only one cbVG junction in each P1 stock, with fewer than 5 reads, compared to 678,462 reads in our SeV Cantell control stock, which has been passaged in conditions to maintain a high content of cbVGs (Fig. 1D). Lastly, we compared the immunostimulatory activity of our stocks to that of the SeV Cantell wild type control. Our newly rescued clean stock (rS1 P1) triggered much lower antiviral gene expression than SeV Cantell and our cbVG-low SeV Cantell stock, which was prepared by growing SeV Cantell under conditions that deplete cbVGs (24), while maintaining similar replication levels (Figs. 1E, F). Together, these data demonstrate that SeV Cantell can be prepared to contain nearly undetectable levels of cbVGs when launched from plasmids.

**Figure S1:**
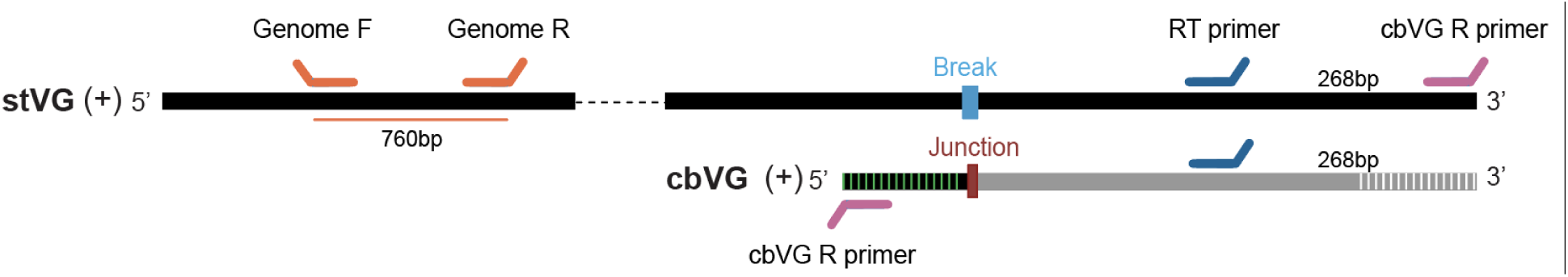
cbVG-PCR schematic. Diagram of the standard viral genome (stVG) and a representative copy-back viral genome (cbVG) of unknown length. The break position of the cbVG is indicated with light blue on the stVG, and the junction rejoin site is marked with burgundy on the cbVG, with the complementary regions represented with black/green stripes and grey/white stripes. The blue RT primer is located 268 bp from the 3’ end of the antigenome. The pink primer represents the reverse primer for cbVG detection, while the orange primers correspond to genomic PCR primers. The orange line below indicates the expected PCR product length of 760 bp.

**Figure 1:**
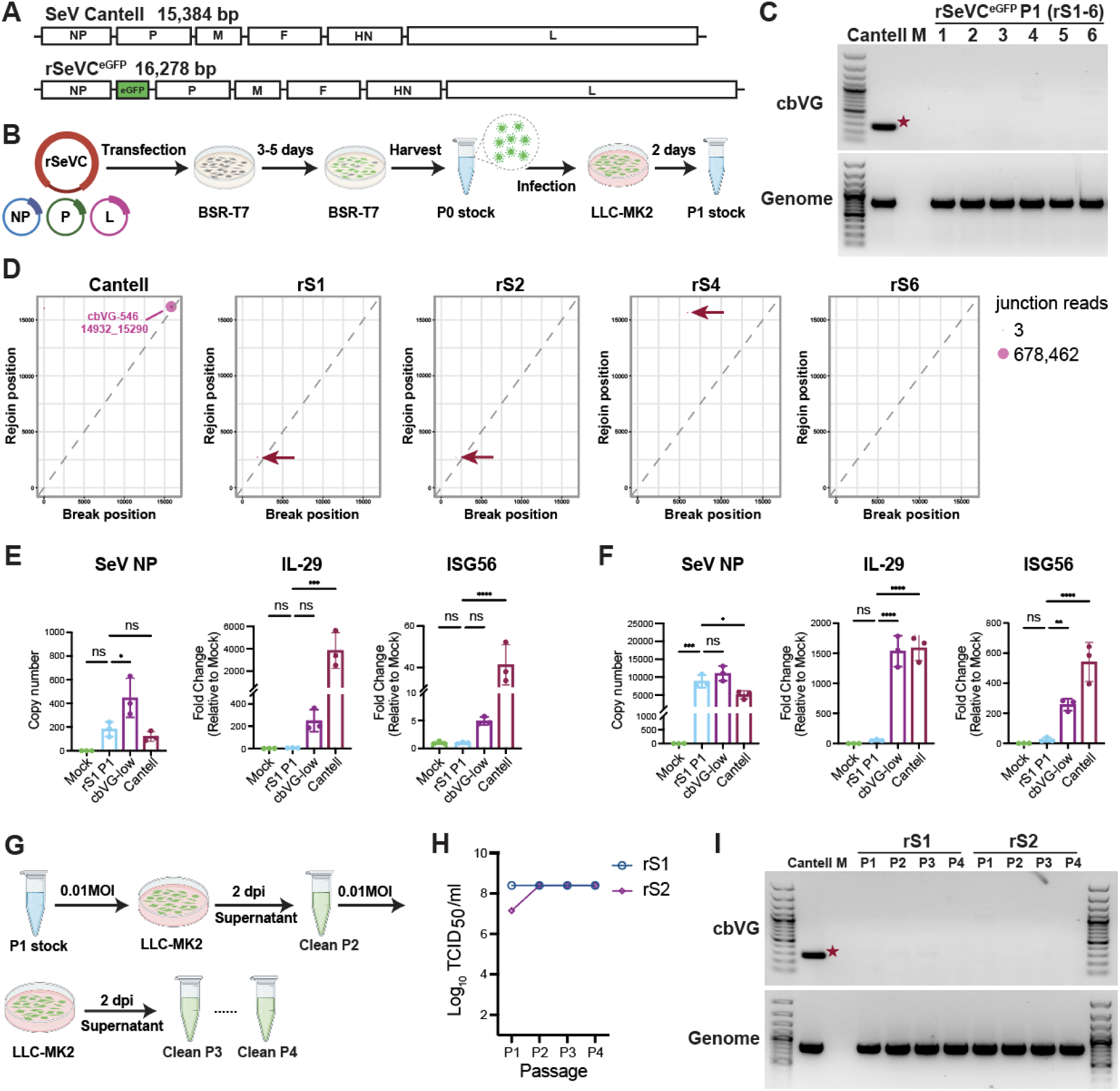
Generation of Sendai virus cbVG-clean stocks using reverse genetics. (A) Schematic diagrams of the SeV genome and the recombinant reporter virus rSeVC^eGFP^. The lengths of the genomes are indicated. (B) Schematic illustration of the SeV rescue process and the development of Passage 1 (P1) clean virus stocks. (C) cbVG and genomic PCR of LLC-MK2 cells infected at an MOI of 1.5 with each of the P1 stocks for 24h. SeV Cantell stock containing high levels of cbVG-546 (indicated by the red star) was used as a positive control. Mock (M) indicates mock-infected cells. (D) Break-rejoin plots displaying cbVG junctions detected by RNA Seq and VODKA2 analysis for SeV Cantell stock and four independently rescued P1 rSeVC^eGFP^ clean virus stocks. Each dot represents a unique cbVG junction, with dot size indicating the read count. The red arrows point to the only cbVG junction present in each clean virus stock. (E, F) SeV NP, IL29 and ISG56 mRNA levels of A549 cells mock treated or infected with rSeVC^eGFP^ clean stocks rS1 P1, SeV Cantell or cbVG-low SeV Cantell stocks at an MOI of 1.5 for 6h (E) or 24h (F). Data represent three independent experiments. Statistical analysis was performed using ordinary one-way ANOVA followed by Dunnett’s multiple comparisons *post hoc* test. ns: not significant, *: p<0.05, **:p<0.01, ***: p<0.001, ****: p<0.0001. (G) Schematic illustration depicting the process of passaging SeV stocks at low MOI from Passage 2 (P2) to Passage 4 (P4). (H) Virus titers from P1 to P4 for the indicated stocks. (I) cbVG and genomic PCR of LLC-MK2 cells infected at an MOI of 0.01 with P1 to P4 of the indicated stocks for 24h. SeV Cantell stock was used as a positive control with cbVG-546 indicated by the red star. Mock (M) indicates uninfected cells.

To evaluate whether the stocks could maintain a clean status over successive passages, we used two stocks, rS1 and rS2, to infect the highly susceptible cell line LLC-MK2 cells at an MOI of 0.01 for 2 days. The viruses were then harvested, and infection was repeated, passaging up to the fourth passage (P4) (Fig. 1G). Throughout these passages, we observed no significant changes in the virus titers for both stocks (Fig. 1H), and no clear cbVG species were detected by PCR, indicating that the clean status of the stocks was maintained through these passages (Fig. 1I).

### Unique populations of cbVG are generated *de novo* from clean virus stocks

To test if high-MOI passaged stocks generated from clean rSeVC^eGFP^ P1 stocks contain the dominant cbVG-546 species present in the original SeV Cantell (20), we generated high-MOI passaged stocks from all our independent rescues. To do this, we passaged rS1, rS2, rS4, and rS6 P1 stocks at a high MOI of 10 (23) until Passage 8 (P8), while rS3 and rS5 were only passaged to P3 (Fig. 2A). RNA sequencing and VODKA2 analyses showed that all P3 and P8 high-MOI passaged stocks contained significantly higher levels of cbVGs compared to clean P1 stocks, although the amount of cbVG varied among different high-MOI passaged stocks. rS1 and rS2 had higher levels of cbVGs than the other four stocks at P3 (Fig. 2B). As expected, the ratio of total cbVG reads to total viral reads was considerably higher in high-MOI passaged stocks than in the clean stocks (Fig. 2C). As cbVGs are known to interfere with standard viral genome replication by competing for the viral polymerase and other replication resources, inducing antiviral immune signaling pathways, and inhibiting viral protein translation (27), we next measured infectious particles production across passages. As expected, we found less infectious viral particles in rS1 and rS2 comparted to rS4 and rS6 at earlier passage (P4) corresponding with the higher amount of cbVGs detected in rS1 and rS2 at P3. In contrast, rS4 and rS6 did not show a significant decrease in infectious particles (Fig. 2D). cbVG PCR of P3 showed that all stocks except rS4 had detectable and different cbVG amplicons, while the cbVGs in rS4 were undetectable (Fig. 2E). None of the P3 stocks showed a dominant amplicon corresponding to cbVG-546, demonstrating that diverse populations of cbVGs are generated *de novo* by SeV Cantell by P3.

**Figure 2:**
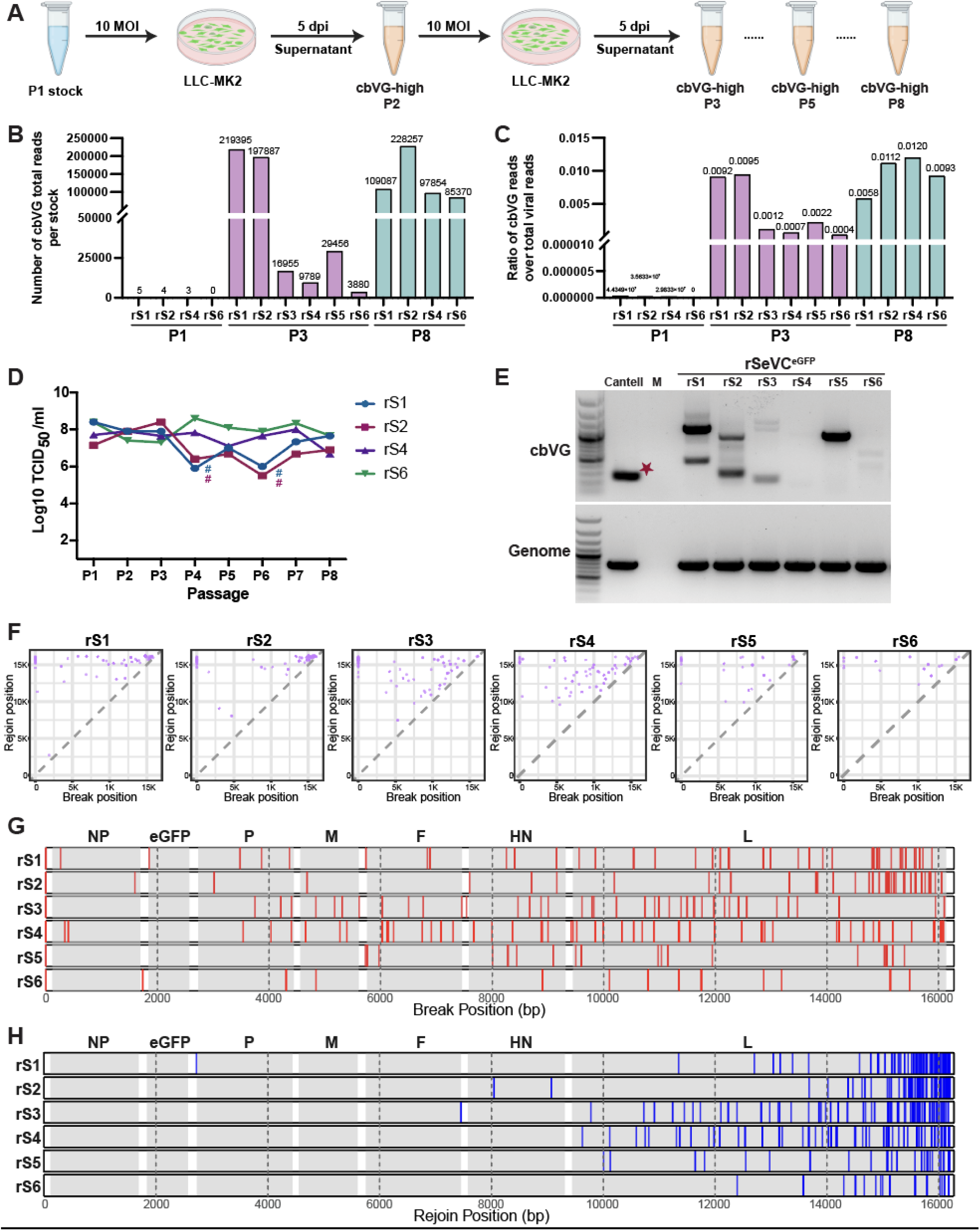
Unique cbVG populations generated by high MOI passaging of clean stocks. (A) Schematic illustration of the process of creating SeV high-MOI passaged stocks from P2 to Passage 8 (P8) starting from P1 clean stocks. (B) Raw number of cbVG reads from total RNA-Seq of clean P1 stocks and high-MOI passaged P3 and P8 stocks. (C) Ratio of total cbVG reads to total viral reads for clean P1 stocks and high-MOI passaged P3 and P8 stocks. (D) Infectious virus titers for rS1, rS2, rS4 and rS6 from P1 to P8. # indicates passages where the titers were too low to conduct an infection at 10 MOI, and an MOI of 0.1 was used instead. (E) cbVG and genomic PCR of LLC-MK2 cells infected at an MOI of 0.1 with each high-MOI passaged P3 virus stock or SeV Cantell for 24h. cbVG-546 was indicated by the red star. Mock (M) indicates uninfected cells. (F) Break-rejoin plots displaying cbVG junctions identified in the six independently high-MOI passaged P3 stocks. Each dot represents a unique cbVG junction. The X-axis and Y-axis represent the break positions and rejoin positions in the parental SeV genome, respectively. (G) Comparison of break positions from (F) across stocks in the genome. Each red line indicates a unique break position. Genes are shaded in gray. (H) Comparison of rejoin positions in the genome from (F) across stocks. Each blue line indicates a unique rejoin position. Genes are shaded in gray.

To further characterize the *de novo*-generated cbVG populations, we identified the cbVGs present in the high-MOI passaged P3 stocks by RNA-Seq. VODKA2 identified populations of cbVGs that significantly differ in each stock, with only few cbVGs with similar break_rejoin positions detected in more than one stock (Fig. 2F). These data corresponded well with the diversity of species detected by PCR (Fig. 2E). The most dominant *de novo-*generated cbVG species in each stock (> 1% of the cbVG population) were formed by interrupting replication (break) throughout the viral genome, including at nucleotide 1 (Fig. 2G), and rejoining near the trailer end of the genome (Fig. 2H). Of note, the most dominant cbVGs in rS1 and rS2 P3 stocks, including those with break at nucleotide 1 of the genome, were recently validated by our group using PCR-independent direct RNA nanopore sequencing (28). Our results agree with observations in the RSV and in measles virus, which showed a large diversity in sequences for the polymerase break, while a much more conserved region was observed for the polymerase rejoin during cbVG generation (22, 29).

### cbVG formation initiates with genome-wide breaks, with position 1 as a conserved feature

To assess the degree of conservation of the positions used for break and rejoin during cbVG generation across the different stocks, we first quantified the number of breaks, rejoins, and junctions (break_rejoin pairs) and compared their occurrences in the high-MOI passaged P3 stocks. We identified 25 break positions, 72 rejoin positions and 25 junctions (break_rejon) that occurred in two or more stocks (Fig. 3A-C). The specific locations and percentages of the 25 recurring break positions in each stock are shown in Fig. 3D with consecutive nucleotides indicated with a horizontal line below the x axis. Of note, all stocks have cbVG breaks at position 1 at a relatively high percentage. The rejoins occurring in two or more stocks showed a distinctive hotspot from positions 16173 to 16203 on the rSeVC^eGFP^ genome (Fig. 3E), which also includes the rejoin position 16185 corresponding to the rejoin of cbVG-546 (position 15290 in the SeV Cantell genome). No similarity in the genomic sequences at the break or rejoin positions was identified (Figure S2). Strikingly, all 25 junctions that were found in 2 or more stocks had a break at the beginning of the genome in positions 1-3 (Fig. 3F), indicating early positions of the genome are frequently rejoined with SeV internal genomic sequences. cbVG-546 was detected at a low percentage (0.0025% among all cbVG reads) in rS2, but not in other P3 stocks. Overall, these analyses demonstrate the limited conservation of cbVG species generated at early virus passage and highlights the diversity of positions for polymerase break and initiation of cbVG formation within the SeV genome, with species breaking at or close to nucleotide 1 as a conserved feature during cbVG formation.

**Figure 3:**
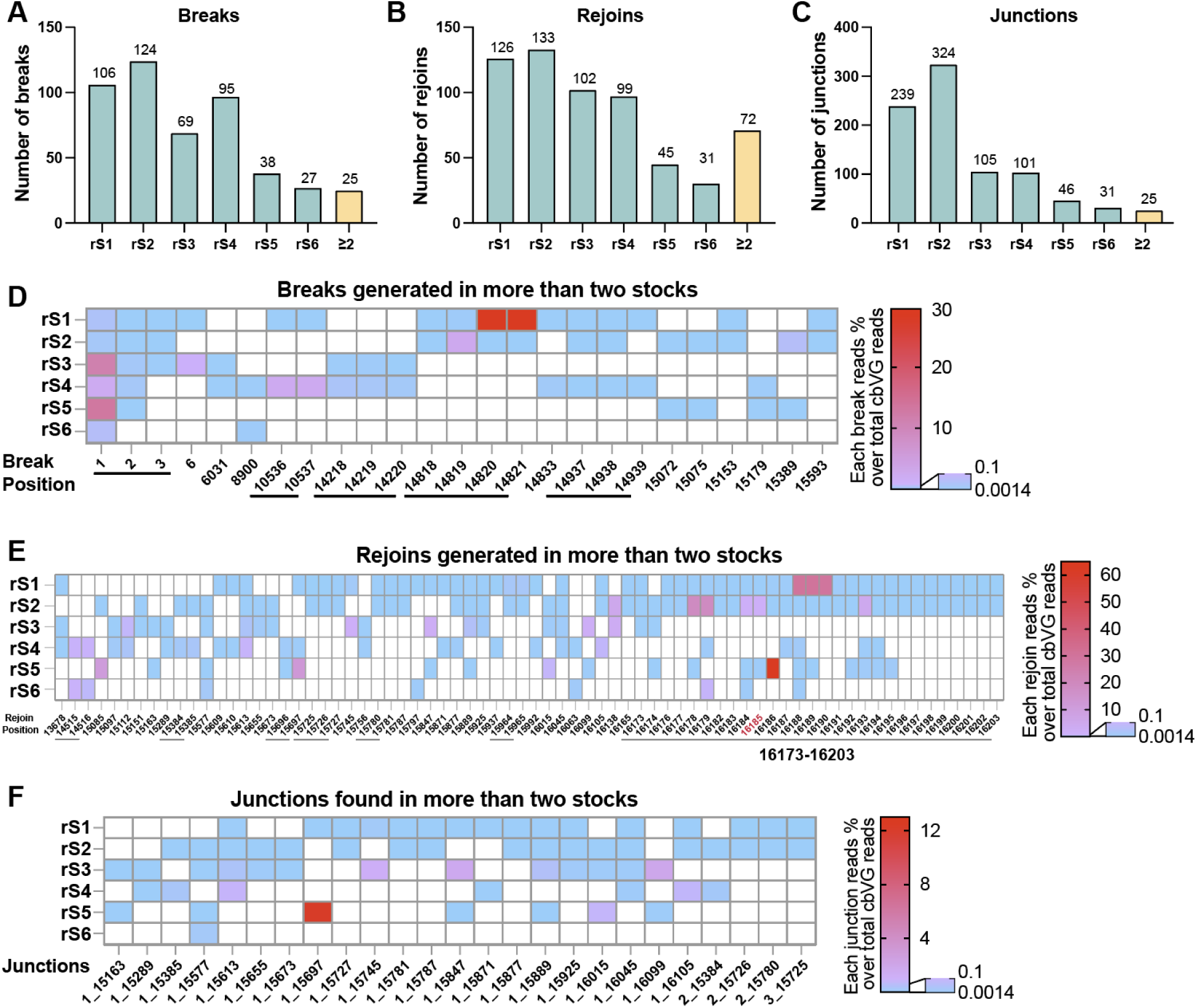
cbVG break, rejoin and junction positions in high-MOI passaged P3 stocks. (A-C) Number of unique breaks (A), rejoins (B) or junctions (break_rejoin) (C) for each high-MOI passaged P3 stock. “≥2” indicates the number of events that appear in two or more stocks. The numbers above the bars indicate the actual number of events. (D-F) Depiction of breaks (D), rejoins (E), and junctions (F) that appear in two or more stocks. The colors indicate the percentages of those breaks, rejoins or junctions over the total cbVG reads in their respective stocks following the corresponding scale on the right. Underlines indicate adjacent break or rejoin positions. The rejoin position in red font in (E) corresponds to the rejoin of cbVG-546 in SeV Cantell.

**Figure S2:**
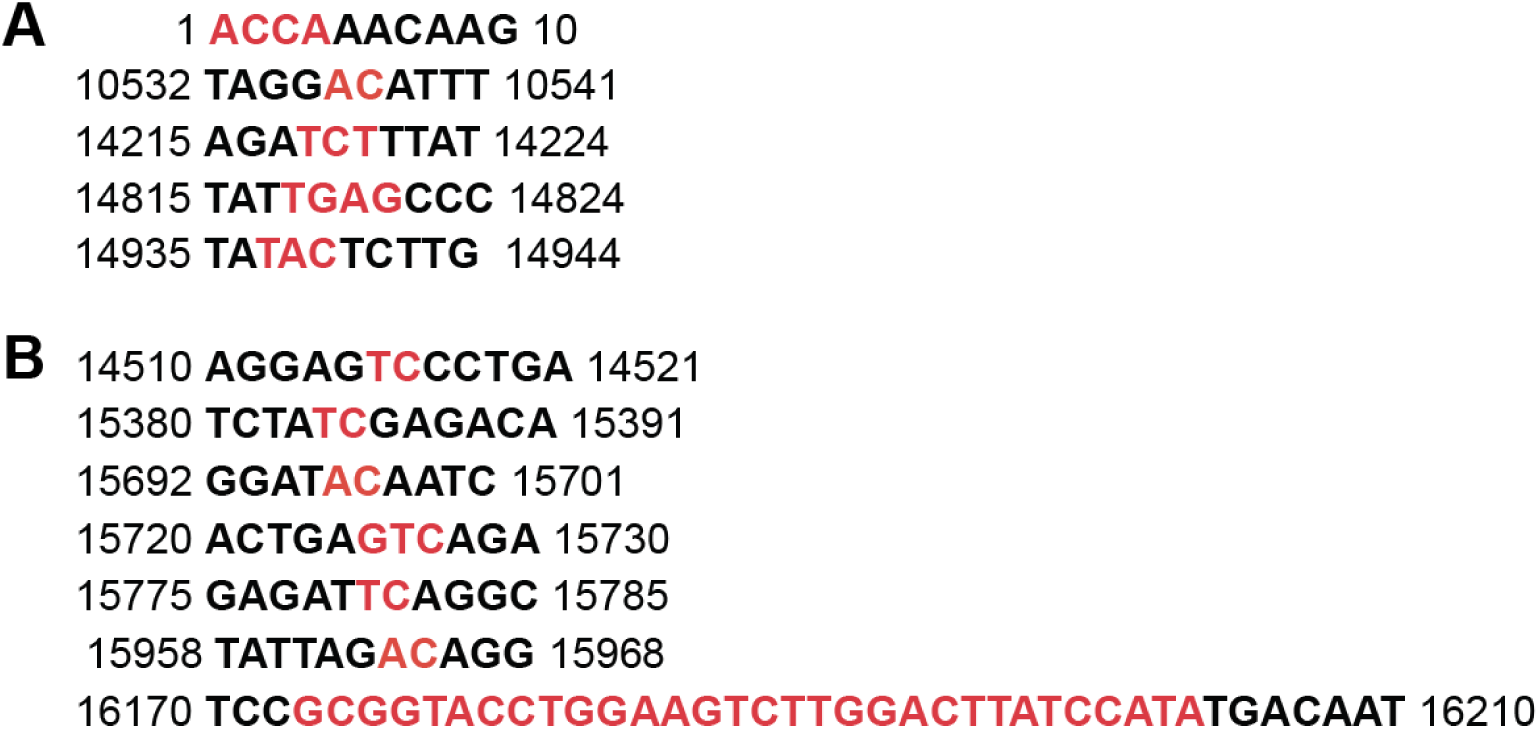
Sequences near adjacent breaks or rejoins. (A) The sequences represent the adjacent breaks and the nearby sequence, with breaks shown in red and the nearby sequence in black. (B) The sequences represent the adjacent rejoins and the nearby sequence, with rejoins shown in red and the nearby sequence in black.

### A subset of cbVGs dominates in each stock independent of the cbVG predicted length

To identify properties that could explain cbVG enrichment during viral passaging, we next analyzed the dominant species in each high-MOI passaged P3 stock. As shown in Fig. S3A, most of the cbVG junctions in P3 were detected at low number of reads, with a raw median read count around 10. In each stock, the number of cbVG reads that reached 1% of the total cbVG reads ranged from 4 to 13 (Fig. 4A). Interestingly, some junctions were highly represented at this early passage, suggesting that these cbVG species rapidly reached high abundance within the population. The dominant cbVGs (>1% of the total cbVG reads) in each stock at this passage varied with no obvious selection of cbVGs of predicted short length (Fig. 4B, 4C and S3B). In fact, rS3 and rS5 had dominant cbVGs with break positions at 1, predicted to be long cbVGs. Dominant cbVG junctions are shown in Table 1. Some of them have similar break and rejoin positions (break_rejoin is ±1 or 2), suggesting that they belong to the same cbVG species.

**Figure 4:**
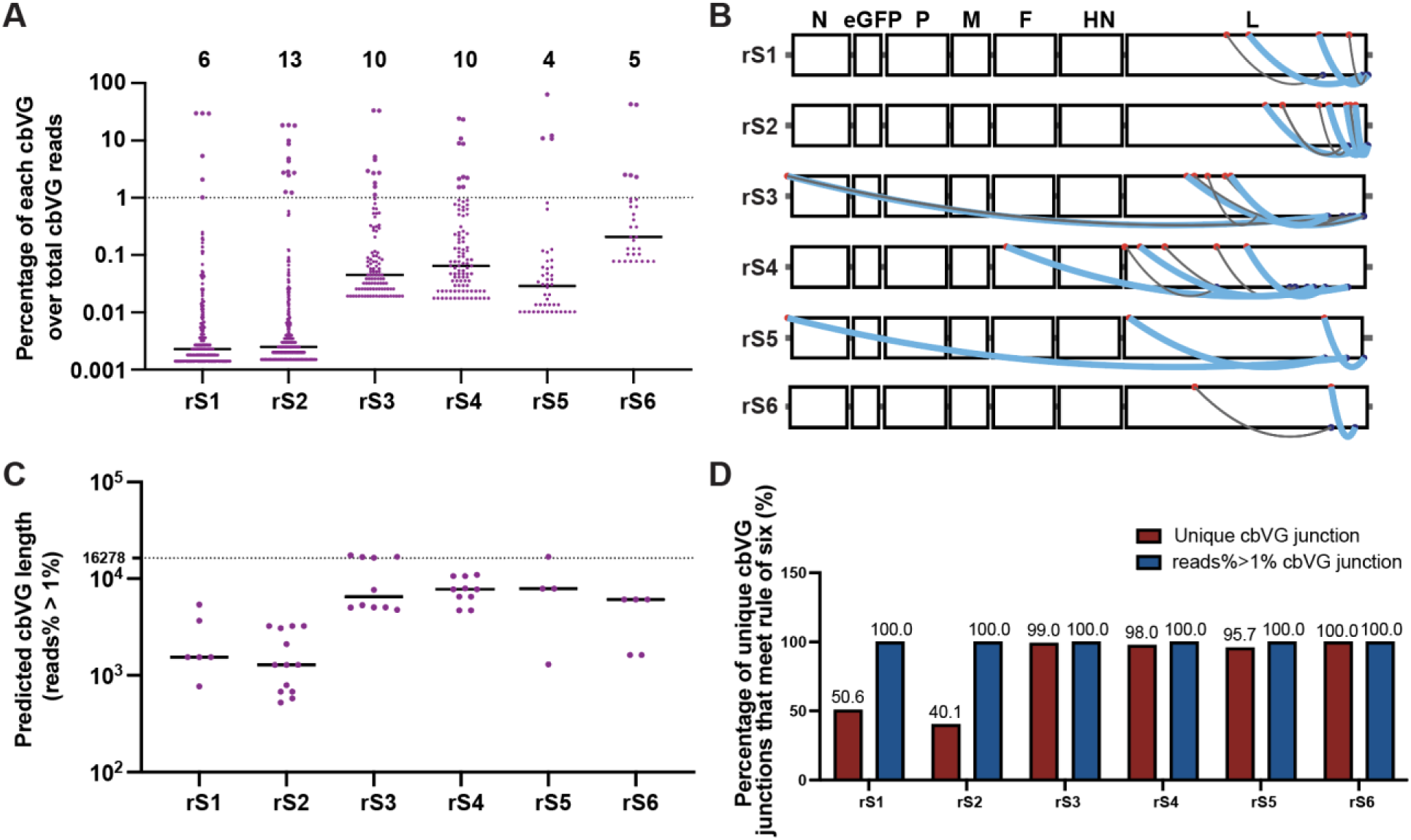
Dominant break positions, rejoin positions, and junctions in each high-MOI passaged viral stock. (A) Percentage of reads for each unique cbVG in each high-MOI passaged P3 stock. Each dot represents a unique cbVG junction, and the black horizontal line indicates the median percentage for each stock. The number on top indicates the number of dominant unique cbVG junctions defined as species with >1% of the total cbVG reads. (B) Mapping of dominant cbVGs from (A) on the rSeVC^eGFP^ genome. Red dots indicate break positions, blue dots indicate rejoin positions. Thick blue lines represent cbVGs with frequencies greater than or equal to 3% of the total cbVG reads. Thin gray lines represent cbVGs with frequencies of 1% to 3% of the total cbVG reads. (C) Dominant cbVGs and their predicted length for all high-MOI passaged P3 stocks. Each dot represents a unique cbVG species. The black horizontal line indicates the median dominant cbVG length for each stock. The dashed line indicates the length of the standard genome. (D) Percentage of unique cbVG junctions (red) and dominant (reads% > 1%) cbVG junctions (blue) in each high-MOI passaged P3 stock that meet the ‘rule of six’.

**Figure S3:**
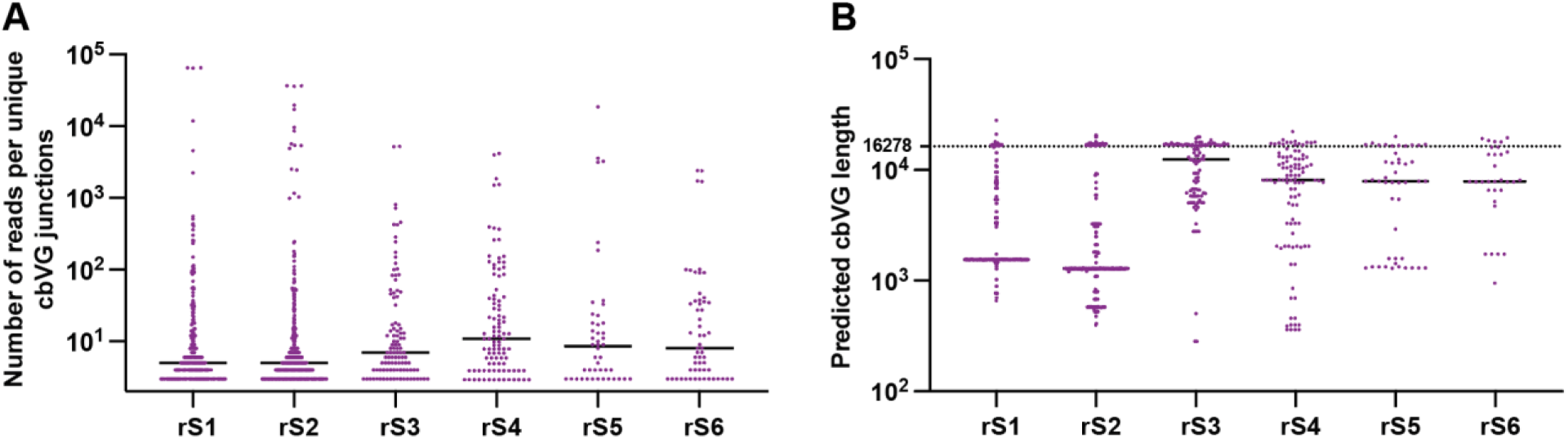
Raw reads and predicted length of all unique cbVGs. (A) Number of reads per unique cbVGs junction for each high-MOI passaged P3 stock. Each dot represents a unique cbVG junction. The black horizontal lines indicate the median number of cbVG reads for each stock. (B) cbVGs and their predicted length for each high-MOI passaged P3 stock. Each dot represents a unique cbVG. The black horizontal lines indicate the median cbVG length for each stock. The dashed line indicates the length of the standard genome.

**Table 1:** Dominant cbVG junctions of each P3 high-MOI passaged stock. * Gray shading indicates cbVGs with adjacent break_rejoin positions that are likely to belong to the same species.

| rS1 |  |  | rS2 |  |  | rS3 |  |  | rS4 |  |  | rS5 |  |  | rS6 |  |  |
| --- | --- | --- | --- | --- | --- | --- | --- | --- | --- | --- | --- | --- | --- | --- | --- | --- | --- |
| Break_Rejoin | Length | % | Break_Rejoin | Length | % | Break_Rejoin | Length | % | Break_Rejoin | Length | % | Break_Rejoin | Length | % | Break_Rejoin | Length | % |
| 14821_16189 | 1548 | 29.6 | 15096_16178 | 1284 | 18.5 | 12412_15100 | 5046 | 33.4 | 12827_15021 | 4710 | 24.1 | 15070_16186 | 1302 | 63.4 | 15137_15795 | 1626 | 43.0 |
| 14822_16188 | 1548 | 29.6 | 15095_16179 | 1284 | 18.4 | 12411_15101 | 5046 | 33.2 | 12826_15022 | 4710 | 23.1 | 1_15697 | 16860 | 12.2 | 15138_15794 | 1626 | 42.2 |
| 14820_16190 | 1548 | 29.3 | 15094_16180 | 1284 | 18.2 | 11174_16062 | 5322 | 5.2 | 9849_14705 | 8004 | 10.8 | 9596_15084 | 7878 | 11.0 | 11350_15130 | 6078 | 2.5 |
| 12858_16016 | 3684 | 5.4 | 13325_16155 | 3078 | 9.9 | 1_15133 | 17424 | 4.6 | 6119_15651 | 10788 | 8.9 | 9595_15085 | 7878 | 10.7 | 11351_15129 | 6078 | 2.5 |
| 15660_16130 | 768 | 2.1 | 15585_16181 | 792 | 8.7 | 11386_16138 | 5034 | 2.9 | 6118_15652 | 10788 | 8.8 |  |  |  | 11349_15131 | 6078 | 2.3 |
| 12238_14926 | 5394 | 1.0 | 15844_16138 | 576 | 4.9 | 12251_15531 | 4776 | 2.7 | 10536_14186 | 7836 | 2.3 |  |  |  |  |  |  |
|  |  |  | 15843_16193 | 522 | 4.3 | 11760_13130 | 7668 | 2.7 | 9417_11987 | 11154 | 2.2 |  |  |  |  |  |  |
|  |  |  | 13803_15515 | 3240 | 2.9 | 1_16099 | 16458 | 1.8 | 10537_14185 | 7836 | 2.1 |  |  |  |  |  |  |
|  |  |  | 13804_15514 | 3240 | 2.7 | 1_15847 | 16710 | 1.6 | 11987_14031 | 6540 | 1.5 |  |  |  |  |  |  |
|  |  |  | 13805_15513 | 3240 | 2.7 | 1_15745 | 16812 | 1.1 | 11986_14032 | 6540 | 1.5 |  |  |  |  |  |  |
|  |  |  | 14819_15633 | 2106 | 2.5 |  |  |  |  |  |  |  |  |  |  |  |  |
|  |  |  | 15696_16184 | 678 | 1.3 |  |  |  |  |  |  |  |  |  |  |  |  |
|  |  |  | 15695_16185 | 678 | 1.2 |  |  |  |  |  |  |  |  |  |  |  |  |

We then assessed whether the dominant cbVGs meet the ‘rule of six’ which establishes that a genome length of a multiple of six is required for optimal paramyxovirus replication (30, 31). As shown in Fig. 4D, 40.1-99.0% of the unique cbVGs met the ‘rule of six’ and all of the dominant cbVGs met this rule, indicating that meeting the rule of six is a critical factor in their selection.

### Dominant cbVGs remain stable through serial passaging, while low abundance cbVGs continue to diversify

To assess how stable the population of cbVGs is after initial selection, we analyzed the cbVGs of rS1, rS2, rS4, and rS6 up to P8. We selected these stocks as representatives of two stocks with a high level of cbVGs (rS1, rS2) and two with a low level of cbVGs (rS4, rS6) based on P3 sequencing and cbVG PCR results. As shown in Figures 5A and 5B, cbVGs were first detected by PCR in P2 in rS1 and rS2, and the dominant species remained stable. However, no clear dominant cbVG bands were seen in rS4 at any passage (Fig. 5C). rS6 showed an intermediate phenotype where amplicons were detected beginning in P3 (Fig. 5D).

**Figure 5:**
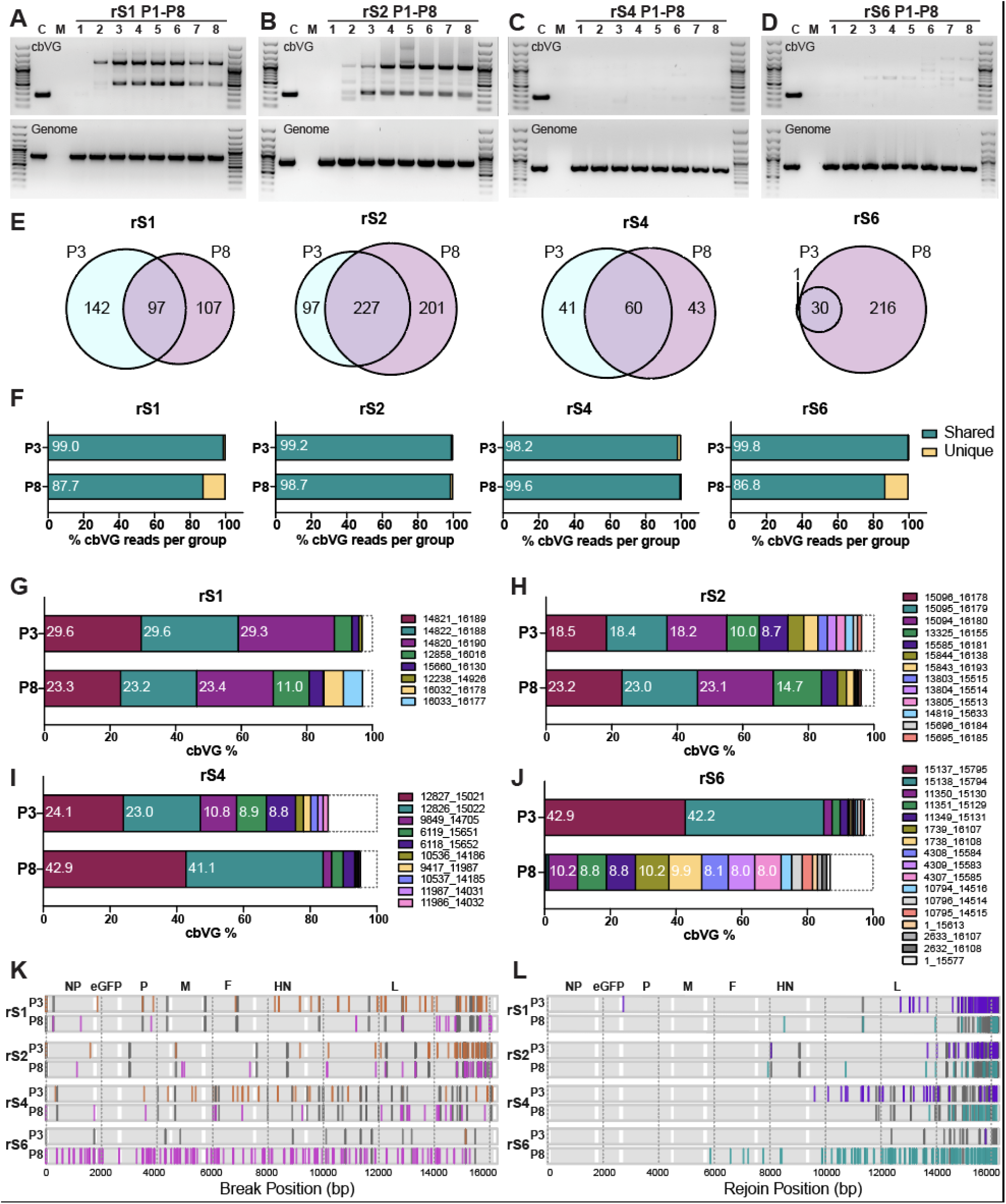
cbVG profiles across passages. (A-D) cbVG and genomic PCR results of high-MOI passaged stocks rS1 (A), rS2 (B), rS4 (C), and rS6 (D) from passages P1 to P8 in LLC-MK2 cells infected at 1.5 MOI for 24h. SeV Cantell (C) containing the predominant cbVG-546 was used as a positive control. Mock (M) indicates uninfected cells. (E) Venn diagrams show the number of unique cbVG junctions in P3 and P8, as well as shared cbVGs between P3 and P8, for each stock. Numbers indicate the cbVG junction counts in each group. (F) Percentage of cbVG reads from shared (teal) and unique (yellow) groups between P3 and P8 for each stock. Numbers inside bars indicate the percentage of shared cbVG reads over total cbVG reads. (G– J) Composition of dominant cbVGs (≥1% of total cbVG reads) in P3 and P8 for rS1 (G), rS2 (H), rS4 (I), and rS6 (J). Dominant cbVGs present in P3, P8, or both are shown. Each color represents a different cbVG junction. Numbers within bars indicate the percentage of each cbVG junction relative to the total cbVG reads (only those ≥8% of total cbVG reads are shown). The dashed box indicates all the other cbVGs. (K) Comparison of break positions between P3 and P8 in the genome. Dark gray lines indicate shared breaks between P3 and P8. Each orange or magenta line indicates a unique break position only in P3 or P8. Genes are shaded in gray. (L) Comparison of rejoin positions between P3 and P8 in the genome. Dark gray lines indicate common rejoins between P3 and P8. Each purple or teal line indicates a unique rejoin position in P3 or P8. Genes are shaded in gray.

We next analyzed RNA-Seq data on P8 stocks of rS1, rS2, rS4, and rS6, and looked for cbVG junctions present only in P3 or P8, or those shared in both passages. Shared unique cbVGs junctions were present in both P3 and P8 in all stocks (Fig. 5E) and the fraction of total shared cbVG reads reached 86.8% - 99.8% among total cbVG reads (Fig. 5F). Although not all dominant cbVGs were shared between P3 and P8, and two dominant cbVGs showed a marked decline in abundance from P3 to P8 in rS6, the most abundant cbVGs in rS1, rS2, and rS4 remained consistent between P3 and P8. (Fig 5G-J). Comparison of breaks and rejoins of the unique cbVGs (those present in only one of the passages) revealed breaks and rejoins that were present in P3 and absent in P8, accompanied by newly generated breaks present only in P8 across the genome (Fig 5K and 5L). These results suggest that in most cases dominant cbVGs remain stable during passaging while low-abundance cbVGs continue to diversify. However, notable changes in the cbVG population were observed in stock rS6, suggesting that cbVG population stability can vary depending on the cbVG population composition.

### cbVG dominant species do not associate with specific SeV genomic variants

Lastly, we assessed whether the generation or accumulation of dominant cbVG species associated with SeV genomic variants that arose during passages. To identify SeV variants, we used the bioinformatics tool iVar (32) that calls variants from sequencing data by aligning reads to a reference genome. As shown in Tables 2–5, a few variants were detected in each stock, with P8 containing more variants than earlier passages, as expected. At P3, only rS1 carries a high-frequency non-synonymous variant (A9996C) in the L protein, resulting in a methionine to leucine substitution. In addition, T20A and T24A variants were detected in the leader. Non-coding region variants were detected in all four stocks, with a higher frequency in P8, indicating that these sites are highly variable. In conclusion, no significant variants were identified that could explain differences in cbVG species or abundance between stocks.

**Table 2:** Variants detected in rS1 stock at Passage 1, 3 and 8.

| Position | Ref | Alt | Alt_freq |  |  | Gene | REF_codon | REF_AA | ALT_codon | ALT_AA |
| --- | --- | --- | --- | --- | --- | --- | --- | --- | --- | --- |
|  |  |  | P1 | P3 | P8 |  |  |  |  |  |
| 20 | T | A |  |  | 0.93 |  |  | / |  | / |
| 24 | T | A |  |  | 0.19 |  |  | / |  | / |
| 24 | T | G |  |  | 0.09 |  |  | / |  | / |
| 58 | G | A |  |  | 0.44 |  |  | / |  | / |
| 274 | T | C |  |  | 0.08 | N | TTC | Phe | TCC | Ser |
| 2999 | A | G | 0.06 | 0.06 |  | P | AGT | Ser | GGT | Gly |
| 3684 | A | +G | 0.07 |  |  | P gene editing |  |  |  |  |
| 4444 | G | A | 0.81 | 0.82 | 0.63 | P | TAG | Stop | TAA | Stop |
| 6273 | G | A |  |  | 0.24 | F | GTG | Val | ATG | Met |
| 9996 | A | C | 0.82 | 0.83 | 0.63 | L | ATG | Met | CTG | Leu |
| 15250 | T | C |  | 0.30 | 0.21 | L | CTT | Leu | CCT | Pro |
| 15560 | T | C | 0.05 |  |  | L | CTT | Leu | CTC | Leu |

\* Gray shading indicates variants with a frequency higher than 0.5.

**Table 3:** Variants detected in rS2 stock at Passage 1, 3 and 8.

| Position | Ref | Alt | Alt_freq |  |  | Gene | REF_codon | REF_AA | ALT_codon | ALT_AA |
| --- | --- | --- | --- | --- | --- | --- | --- | --- | --- | --- |
|  |  |  | P1 | P3 | P8 |  |  |  |  |  |
| 20 | T | A |  |  | 0.92 |  |  | / |  | / |
| 24 | T | A |  |  | 0.82 |  |  | / |  | / |
| 58 | G | A |  |  | 0.07 |  |  | / |  | / |
| 3684 | A | +G | 0.08 |  |  | P gene editing |  |  |  |  |
| 7864 | A | G |  |  | 0.65 | HN | CAA | Gln | CGA | Arg |
| 8353 | A | G |  |  | 0.09 | HN | AAA | Lys | AGA | Arg |
| 9325 | C | A |  |  | 0.08 | Between HN/L |  |  |  |  |
| 15719 | A | T | 0.16 | 0.16 | 0.23 | L | ATA | Ile | ATT | Ile |

\* Gray shading indicates variants with a frequency higher than 0.5.

**Table 4:** Variants detected in rS4 stock at Passage 1, 3 and 8.

| Position | Ref | Alt | Alt_freq |  |  | Gene | REF_codon | REF_AA | ALT_codon | ALT_AA |
| --- | --- | --- | --- | --- | --- | --- | --- | --- | --- | --- |
|  |  |  | P1 | P3 | P8 |  |  |  |  |  |
| 20 | T | A |  | 0.19 | 0.72 |  |  | / |  | / |
| 24 | T | A |  | 0.09 | 0.75 |  |  | / |  | / |
| 58 | G | A |  | 0.05 |  |  |  | / |  | / |
| 2450 | A | C | 0.08 | 0.09 | 0.08 | eGFP | AGC | Ser | CGC | Arg |
| 2451 | G | A | 0.08 | 0.09 | 0.08 | eGFP | AGC | Ser | AAC | Asn |
| 3221 | C | T | 0.28 | 0.27 | 0.11 | P | CCT | Pro | TCT | Ser |
| 3684 | A | +G |  | 0.06 | 0.10 | P gene editing |  |  |  |  |
| 4094 | A | G |  |  | 0.07 | P | AAA | Lys | GAA | Glu |
| 6167 | C | T | 0.05 |  |  | F | ACC | Thr | ACT | Thr |
| 7372 | T | C | 0.06 | 0.06 | 0.15 | F | ATA | Ile | ACA | Thr |
| 7497 | A | T |  | 0.07 | 0.21 | Between F/HN |  | / |  | / |
| 7582 | A | C | 0.24 | 0.21 | 0.07 | Between F/HN |  | / |  | / |
| 8236 | C | T |  | 0.11 | 0.52 | HN | GCT | Ala | GTT | Val |
| 11128 | A | G |  |  | 0.07 | L | AAG | Lys | AGG | Arg |
| 12825 | A | G |  |  | 0.18 | L | ATC | Ile | GTC | Val |
| 13672 | A | G |  |  | 0.08 | L | GAT | Asp | GGT | Gly |

\* Gray shading indicates variants with a frequency higher than 0.5.

**Table 5:** Variants detected in rS6 stock at Passage 1, 3 and 8.

| Position | Ref | Alt | Alt_freq |  |  | Gene | REF_codon | REF_AA | ALT_codon | ALT_AA |
| --- | --- | --- | --- | --- | --- | --- | --- | --- | --- | --- |
|  |  |  | P1 | P3 | P8 |  |  |  |  |  |
| 20 | T | A |  | 0.19 | 0.46 |  |  | / |  | / |
| 24 | T | A |  | 0.08 | 0.49 |  |  | / |  | / |
| 449 | T | C |  |  | 0.07 | N | CCT | Pro | CCC | Pro |
| 2479 | C | G | 0.19 | 0.17 | 0.08 | eGFP | CCC | Pro | CCG | Pro |
| 3684 | A | +G |  | 0.06 |  | P gene editing |  |  |  |  |

## Discussion

In this study, we address a major challenge in cbVG research: the inability to distinguish between *de novo* cbVG generation and carryover from earlier virus stocks. By generating cbVG-free recombinant Sendai viruses and passaging them under defined conditions, we demonstrate that cbVGs arise reproducibly but with stock-specific compositions, indicating both conserved and stochastic features of their formation. These results advance our understanding of cbVG biology and establish an experimental system to study the mechanisms underlying their generation and accumulation.

Our six high-MOI passaged stocks, generated from independently rescued rSeVC^eGFP^, exhibited distinct cbVG profiles and levels with few conserved features. In the past, the lack of truly cbVG-clean viral stocks limited studies of cbVG generation to identifying the cbVGs present in virus populations and tracking the accumulation of specific cbVGs during serial passaging (33, 34). By contrast, the study of newly rescued recombinant viruses overcomes this limitation, enabling a more systematic characterization of cbVG formation. Notably, studies examining the role of the SeV and measles virus C protein in cbVG generation also revealed the presence of *de novo* generated cbVG species in newly rescued viruses (20, 22); although cbVGs were not the main focus of those studies, their findings support our observation that cbVG generation is highly variable yet retains certain conserved features.

In this study, we observed cbVGs with junctions of position 1, or close to it, and a sequence closer to the trailer of the genome across all six rescued virus stocks, indicating that this species is a frequent occurrence. Similar potentially long cbVGs were detected in RSV-infected *in vitro* samples, as well as in clinical samples but remain understudied (35). Moreover, our recent work using direct RNA sequencing of rSeVC^eGFP^ stocks has further validated their existence (28). Although long cbVGs have often been debated as sequencing or assembly artifacts, our findings provide direct evidence that they are generated during infection and may represent a conserved feature of paramyxovirus replication. Importantly, the presence of these predicted long cbVGs raises the possibility that they play a role in modulating the generation of shorter cbVGs, viral replication dynamics and/or host antiviral responses, warranting further investigation.

While short cbVGs were previously thought to preferentially accumulate as dominant species due to their shorter replication cycles, our results demonstrate that cbVG length is not necessarily linked to dominance. Instead, cbVGs that conform to the ‘rule of six’ showed a clear replication advantage, and all dominant cbVGs in our study adhered to this rule, consistent with previous reports (30). As our analysis was limited to eight passages, whether extended passaging would alter cbVG dynamics remains an open question for future studies.

Comparison of cbVG populations in P3 and P8 stocks showed that new cbVGs continued to arise even in the presence of dominant cbVGs, but they did not accumulate to high levels by passage 8. Together with the persistence of cbVG-546 in the SeV Cantell strain, these findings suggest that once established, dominant cbVGs are difficult to replace, likely due to a combination of replication dynamics, host selection pressures, and potential functional advantages. Although many novel cbVGs were generated in our study, we did not investigate whether they share functional properties such as activating innate immune responses, interfering with genome replication and protein expression, or contributing to persistence. If such functions are conserved, they may be driven by common sequence or structural features. For example, our recent work demonstrated that specific nucleotides in the terminal loop of the cbVGs are necessary for strong activation of RIG-I signaling in cells (36), raising the possibility that similar motifs underlie the activity of newly generated cbVGs.

Since a virus is a community of particles containing different standard and non-standard viral genomes (1), our stocks contained not only cbVGs but also variants of the standard virus, with higher frequencies of variants observed at later passages, as expected. None of the variants could explain the frequency or characteristics of the different cbVG populations. Among the six P3 stocks, two (rS1 and rS2) had high cbVG levels; however, only rS1 carried a high-frequency mutation in the polymerase L gene, indicating that this mutation is not directly linked to cbVG abundance. All stocks had variants at positions 20 and 24 of the leader region, which was more frequent at P8 than at P3, suggesting that this site represents a high-frequency adaptive mutation in LLC-MK2 cells. Previous studies on SeV reported that a single G576T mutation (N protein D153Y) is associated with cbVG production, likely by weakening N-N interactions and reducing nucleocapsid density (23), however, we did not detect this mutation in our stocks, highlighting that the variant landscape may depend on both passage history and host cell context. A recent study on RSV found that a certain cbVG rejoin hotspot is linked to the trailer sequence. When a poly-U mutation was introduced into the trailer, cbVG generation was reduced, but during passaging a viral variant arose that restored cbVG production (37). In our study, however, we did not observe any mutations in the trailer sequence related to cbVG abundance. This difference may be virus-specific. Such arising variants could influence cbVG generation or viral adaptation in ways that extend beyond replication efficiency alone, an area that warrants further investigation.

In summary, by establishing cbVG-free SeV stocks, we were able to directly examine the *de novo* generation and accumulation of cbVGs. Our findings demonstrate that cbVG generation is both frequent and stochastic; yet certain features, such as adherence to the ‘rule of six’ and breakpoints near the position 1 of the genome, appear conserved. These insights highlight the dual nature of cbVG biology, shaped by both random polymerase errors and selective forces that favor cbVG species. Future studies should investigate how cbVG diversity translates into functional outcomes, including their roles in modulating viral replication, shaping host immune responses, and influencing infection persistence. A deeper understanding of these processes will not only clarify the role of cbVGs in viral pathogenesis but may also open new avenues for harnessing cbVGs as antiviral or immunomodulatory tools.

## Materials & methods

### Cell lines

BSR-T7/5 cells (kindly provided by Dr. Conzelmann) (38), LLC-MK2 cells (ATCC, #CCL-7), and A549 cells (ATCC, #CCL-185) were cultured in tissue culture medium (Dubelcco’s modified Eagle’s medium (DMEM) (Invitrogen,#11965092) supplemented with 10% fetal bovine serum (FBS) (Sigma, #F0926), gentamicin 50ng/ml (ThermoFisher, #15750060), L-glutamine 2 mM (Invitrogen, #G7513) and sodium pyruvate 1 mM (Invitrogen, #25-000-C1) at 5% CO2 37°C. Cells were treated with myco-plasma removal agent (MP Biomedical, #3050044) and tested monthly for mycoplasma contamination using the MycoAlert Plus mycoplasma testing kit (Lonza, #LT07-318).

### Virus rescue and passage

Six independent rSeVC^eGFP^ viruses, named rS1 to rS6, were rescued as previously described (25) and tittered as described in dx.doi.org/10.17504/protocols.io.ewov1d73yvr2/v1. high-MOI passaged virus stocks were developed from the rescued viruses as described in dx.doi.org/10.17504/protocols.io.j8nlk9xqwv5r/v1 (28). rS1 and rS2 have been reported previously as rSeVB and rSeVA (28).

### cbVG-PCR

To detect cbVGs by PCR, LLC-MK2 cells were infected with SeV at 1.5 MOI for 24 hours. Total RNA of infected cells and mock cells were extracted using Kingfisher and a Mag-MAX mirVana Total RNA Isolation Kit (Thermo Fisher, #A27828) following manufacturer’s guideline. 250ng of total RNA was used for cDNA synthesis with high-capacity RNA to cDNA kit (Thermo Fisher, #18080051) with our RT primer (24) to capture viral RNA including cbVGs. SeV cbVG RT-PCR was done as described in dx.doi.org/10.17504/protocols.io.5qpvok3j7l4o/v1 (24).

### RT-qPCR

Total RNA from infected cells and control samples was extracted and transcribed as described above. qPCR was performed on an Applied Biosystems QuantStudio 5 system using SYBR Green (Thermo Fisher, #S7564) and 5 μM forward and reverse primers specific for SeV NP, IL29, and ISG56. Primer sequences are listed in Table 6. Relative transcript levels were normalized to human GAPDH and β-actin expression as previously described (39).

**Table 6.**
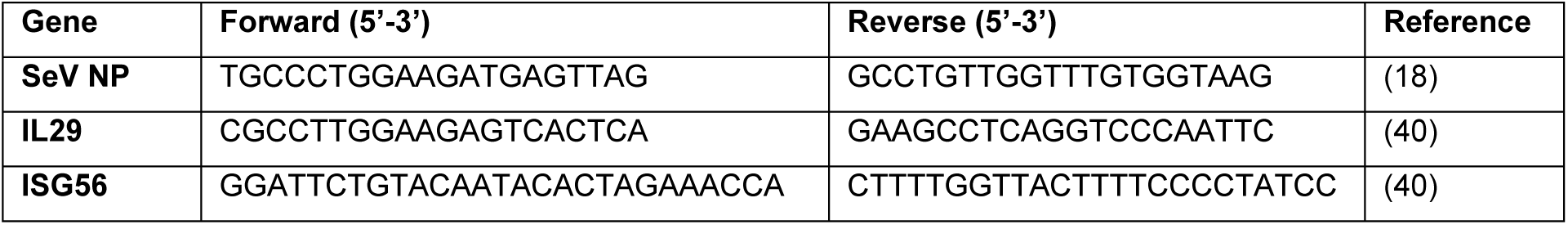
Primers for qPCR.

### RNA sequencing, VODKA2 and iVar

RNA sequencing libraries were prepared using the Illumina TruSeq Stranded Total RNA kit and sequenced on a NextSeq platform (High Output, 2 × 150 bp), generating ∼30 million reads per sample. The raw sequencing data have been deposited in the NCBI Sequence Read Archive (SRA) under the submission number SUB15714354. Data preprocessing and cbVG detection were performed with VODKA2 as previously described (26). In some cases, consecutive nucleotides were identified by VODKA2 as break or rejoin sites. These may represent different cbVGs or a single cbVG that was split into multiple ones during library preparation, sequencing, or bioinformatic processing. For this analysis, we considered them as individual break or rejoin sites. Viral variants were detected using iVar (32).

### Statistics

Statistics were calculated using GraphPad Prism Version 10 (GraphPad Software, San Diego,CA).

## Acknowledgments

We thank Dr. Karl-Klaus Conzelmann for providing the BSR-T7/5 cells, Sydney Faber for library preparation and Emna Achouri for VODKA2 analysis. This work was funded by NIH/NIAID grants A137062, AI188900 and BJC Investigator fund to CBL, and the Alexander & Gertrude Berg fellowship to YY.

